# In Vitro Inactivation of Human Coronavirus by Titania Nanoparticle Coatings and UVC Radiation: Throwing Light on SARS-CoV-2

**DOI:** 10.1101/2020.08.25.265223

**Authors:** Svetlana Khaiboullina, Timsy Uppal, Nikhil Dhabarde, Vaidyanathan Ravi Subramanian, Subhash C. Verma

## Abstract

The newly identified pathogenic human coronavirus, SARS-CoV-2, led to an atypical pneumonia-like severe acute respiratory syndrome (SARS) outbreak called <u>co</u>rona<u>vi</u>rus <u>d</u>isease 2019 (COVID-19). Currently, nearly 23 million cases have been confirmed worldwide with the highest COVID-19 cases been confirmed in the United States. As there is no vaccine or any effective interventions, massive efforts to create a postential vaccine to combat COVID-19 is underway. In the meantime, safety precautions and effective disease control strategies appear to be vital for preventing the virus spread in the public places. Due to the longevity of the virus on smooth surfaces, photocatalytic properties of self-disinfecting/cleaning surfaces appear to be a promising tool to help guide disinfection policies to control infectious SAR-CoV-2 spread in high-traffic areas such as hospitals, grocery stores, airports, schools, and stadiums. Here, we explored the photocatalytic properties of nanosized TiO_2_ (TNPs) as induced by the UV radiation, towards virus deactivation. Our preliminary results using close genetic relative of SAR-CoV-2, HCoV-NL63, showed the virucidal efficacy of photoactive TNPs deposited on glass coverslips, as examined by quantitative RT-PCR and virus culture assays. Efforts to extrapolate the underlying concepts described in this study to SARS-CoV-2 are currently underway.

## 1. Introduction

The novel coronavirus SARS-CoV-2 (2019-nCoV), causative agent of the pneumonia-like COVID-19 pandemic, has infected more than 23 million people globally [1]. Due to extremely transmissible nature of COVID-19 and limited available intensive care unit (ICU) management, over 800,000 people have lost their life to this fatal SARS-CoV-2 outbreak, worldwide [2]. The highest rate of mortality is found among individuals aged 65 and over [3], whereas the death rate continues to decrease, but still a concern among persons aged 85 years and over [4,5]. Devastating consequences of COVID-19 infection and lack of a vaccine or effective antivirals against SARS-CoV-2 warrants scientific research on this newly identified coronavirus for developing therapeutics and interventions to control viral spread.

One of the main reasons for the observed surge in COVID-19 cases could be attributed to multiple modes of virus dissemination, including direct contact with infected person through respiratory droplets generated during coughing, sneezing, or talking, and indirect contact with contaminated surfaces or objects used by infected persons [6,7]. Analysis of 1070 specimens from multiple sites of 205 COVID-19-infected patients revealed the presence of viral load in urine, nasal, blood, sputum, and fecal samples [8]. Additionally, stability of SARS-CoV-2 in the environment also contributes to its spread [7]. Supporting the assumption of non-aerosol virus transmission are documented COVID-19 cases, which could be explained by direct contact [9]. Due to the risk of nosocomial spread, clinical management of critically ill patients and infection prevention among health-care workers and non-COVID-19-infected patients requires the continuous use of personal protection and decontamination practices [10]. In addition, World Health Organization (WHO), recommends that maintaining physical distancing, frequent hand-washing and cleaning of the contaminated surfaces reduces the chance of contracting COVID-19 infection [11].

COVID-19 pandemic brought the importance of environmental hygiene and cleaning practices to prevent viral spread. COVID-19 virus was detected on multiple surfaces in the near proximity to patient’s bed as well as in the sink and the toilet bowl [12]. Investigation of the hospital cluster of cases identified respiratory droplets as the main source of the significant environmental contamination [13]. In another study, transmission of the virus via fomites (e.g. elevator buttons and public restrooms) was suggested in the cluster of COVID-19 cases in a mall in Wenzhou, China [14] as analyzed by real time RT-PCR. However, detection of viral RNA in the environmental samples based on PCR-based arrays may not always indicate the presence of viable virus that could be transmitted. It is especially important to evaluate the pathogen survival in the environment as the COVID-19 virus was shown to retain the infectivity for up to 72 hours on the stainless steel and plastic surfaces, commonly used in the medical settings [7].

The best way to control the spread of the COVID-19 pandemic is to reduce the viral spread. Strategies to control the spread include use of hand-sanitizers, facemasks, physical distancing, self-isolation of patients, regular cleaning of shared surfaces, and frequent use of disinfectants. Since fomites are one of the ways of virus transmission, it is crucial to identify methods that could reduce virus survival on the surfaces. In this regard, generation of self-disinfecting surfaces based on the advanced oxidation processes/photocatalysis initiated by ultraviolet (UV) radiation will be of great importance. UV irradiation-based methods are powerful disinfectants due to its germicidal ability and have shown effectiveness in reducing infections caused by pathogens. Additionally, nanosized titanium dioxide (TiO_2_-nanoparticles or TNPs) have been proven to act as effective photocatalysts and shown to have both bactericidal and virucidal properties [15]. The anti-viral effect of TiO_2_ has been demonstrated against influenza virus [16], which is transmitted via aerosol and causes respiratory tract infection, similar to COVID-19 [17]. However, the anti-viral efficacy of TiO_2_ against human coronavirus remains largely unknown.

In this study, we sought to determine the disinfection ability of TNPs using an in vitro approach. Our data demonstrate that the photocatalytic properties of TNPs are highly effective in inactivating human coronavirus, HCoV-NL63 by reducing the viral genomic RNA stability and virus infectivity. HCoV-NL63 is an alpha coronavirus, which causes acute respiratory distress symptoms (ARDS) among infected individuals [18]. This virus is a Bio Safety Level 2 pathogen and has high similarity with SARS-CoV-2, therefore was used as a surrogate to determine the efficacy of TNPs in virus disinfection [19]. Our results demonstrated that TNP coatings drastically increase the efficacy of HCoV-NL63 inactivation and even a very brief exposure of UV can effectively eliminate infectious virus or viral genomic RNA. Additionally, the efficacy of TNP coatings, in inactivating the virus, was retained at multiple humid environments, suggesting this to be an effective measure for providing clean/disinfected surfaces in public and hospital settings. In conclusion, this study provides conclusive evidence that surfaces coated with TNPs, a non-toxic thin layer applied as paint, can enhances surface disinfection from human coronaviruses.

## 2. Materials and Methods

### 2.1. Cells

Vero E6 and HEK293L cells were maintained in Dulbecco’s modified Eagle medium (DMEM) supplemented with 10% fetal bovine serum (FBS, Atlanta Biologicals), 2 mM L-glutamine, 25 U/mL penicillin, and 25 μg/mL streptomycin. Cells were grown at 37°C in a humidified chamber supplemented with 5% CO_2_.

### 2.2. Human coronavirus

HCoV-NL63 strain, is a human coronavirus and belongs to the family of alpha coronaviruses. HCoV-NL63 was obtained from BEI Resources (1.6×10^6^ TCID_50_/ml; lot 70033870, NIAID, NIH) and propagated in Vero cells by infecting the Vero cell monolayer with HCoV-NL63 for 2 h. Unattached virus was removed by washing followed by addition of fresh medium. After seven days, virus was harvested, cell debris removed by centrifugation, and virus was quantified by quantitative real-time PCR (qRT-PCR). All the assays were conducted under the biosafety level 2+ (BSL-2+) containment.

### 2.3. TiO_2_ nanoparticles

(TNPs)-coated glass coverslips. TiO_2_ was a generous gift from Degussa Corporation (Degussa Corp. Piscataway, NJ). TNPs suspension (300 μg/mL) was prepared in deionized (DI) water and vortexed. An aliquot (600 μL) of TNPs was placed on the clean UV-treated glass coverslip and dried at 60°C, which resulted into a semi-transparent coating. TNPs-coated coverslips were stored at room temperature (RT) until further use.

### 2.4. UV Photocatalysis

An aliquot of HCoV-NL63 virus (100 μL, Median Tissue Culture Infectious dose (TCID_50_) was placed on TNPs-coated and uncoated coverslips (18 mm diameter, 1017.88 mm^2^) and exposed to the USHIO Germicidal Lamp (model G30T8), which generates UV-C light (wavelength: 254nm, 99V, 30W, 0.355A) for various time points. The UV light source was placed 76 cm, 50 cm and 10 cm from the bottom of the wells containing coverslips, which applied calculated 2900 μW/cm^2^, 4300 μW/cm^2^ and 13000 μW/cm^2^ energies, respectively (where μW = 10^-6^J/s). Virus inactivation was also analyzed under different humidity environments, 45%, 65% and 85% relative humidity, by applying a known amount of virus on TNPs coated and uncoated cover-slips placed under indicated humidity. The humidity conditions were generated by using humidifier placed inside the biological safety cabinet (BSL2) and the humidity was measured using AcuRite Indoor Thermometer and Hygrometer with Humidity Gauge (Acurite, Inc.). Humidity range (45-85%) was selected based on the average (45%) standard in-door humidity recommended by Center for Disease control and Prevention (CDC) and the outdoor humidity registered in many parts of the world including coastal areas, which can have humidity as high as high as 85% [20] [21]. Virus from the TNP coated surfaces were recovered and used for total RNA extraction or applied onto a permissive, human embryonic kidney (HEK293L) cell monolayer for the detection of infectious virus. To ensure the efficacy of the virions recovery from TNP surfaces, cover slips were incubated with PBS at 37°C for 30min (3x) and combined before using for infection assays. Total RNA was extracted by adding Trizol reagent directly onto the coverslip to collect RNA from all the viruses.

### 2.5. Virus Infectivity assay

HEK293L cell monolayer was inoculated with HCoV-NL63 virus aliquot for 2 h (37°C, 5% CO_2_). Cells were washed to remove the unbound virus followed by addition of fresh medium. 48 h post-infection, monolayers were harvested adding Tizol reagent onto the cells and used for total RNA extraction.

### 2.6. RNA extraction and qPCR

Total RNA was extracted using Trizol reagent (Invitrogen, Carlsbad, CA) according to the manufacturer’s recommendation. An aliquot of RNA (1 μg) was used for synthesizing the cDNA (Superscript kit; Invitrogen, Carlsbad, CA). cDNA (2μL) was used for the relative quantification of viral genome RNA in a qPCR assay (ThermoFisher Scientific, Waltham, MA). Primers used in this study are NL63-SF 5’-GTGCCATGACCGCTGTTAAT-3’ and NL63-SR 5’-GCGGACGAACAGGAATCAAA-3’. For generating a standard curve to estimate the viral copies, 100 μl of NL63 (BEI resources) stock with known amounts of HCoV-NL63 was used for total RNA extraction and cDNA synthesis. An aliquot (2μl) of HCoV-NL63 cDNA used for qRT-PCR was equivalent to 320 copies of HCoV-NL63 (1.6×10^6^ TCID_50_/ml; lot 70033870, NIAID, NIH). Several ten-fold dilutions of this isolated genomic RNA (BEI Resources) was used for generating a standard curve. The sensitivity of qPCR assay was 3.2 virions in the sample based on calculations using RNA extracted from the BEI NL63 virus stock.

### 2.7. Experimental setup

In order to test the efficacy of TNPs in accelerating the inactivation of HCoV-NL63 upon exposure to UV-light, a suspension of TNPs was prepared and deposited on glass coverslips as a semitransparent film. TNP-coated coverslips were then placed in the center of a 60 mm culture dish and an aliquot of HCoV-NL63 virus was applied on the top of coverslips, followed by exposure to UV emitting lamps for the selected time duration. After TNP/UV-treatment, the inactivated virus was collected and evaluated for the viral genomic RNA stability and virus infectivity. The experimental plan depicting these steps is presented as a graphical image (Figure. 1).

**Figure 1.**
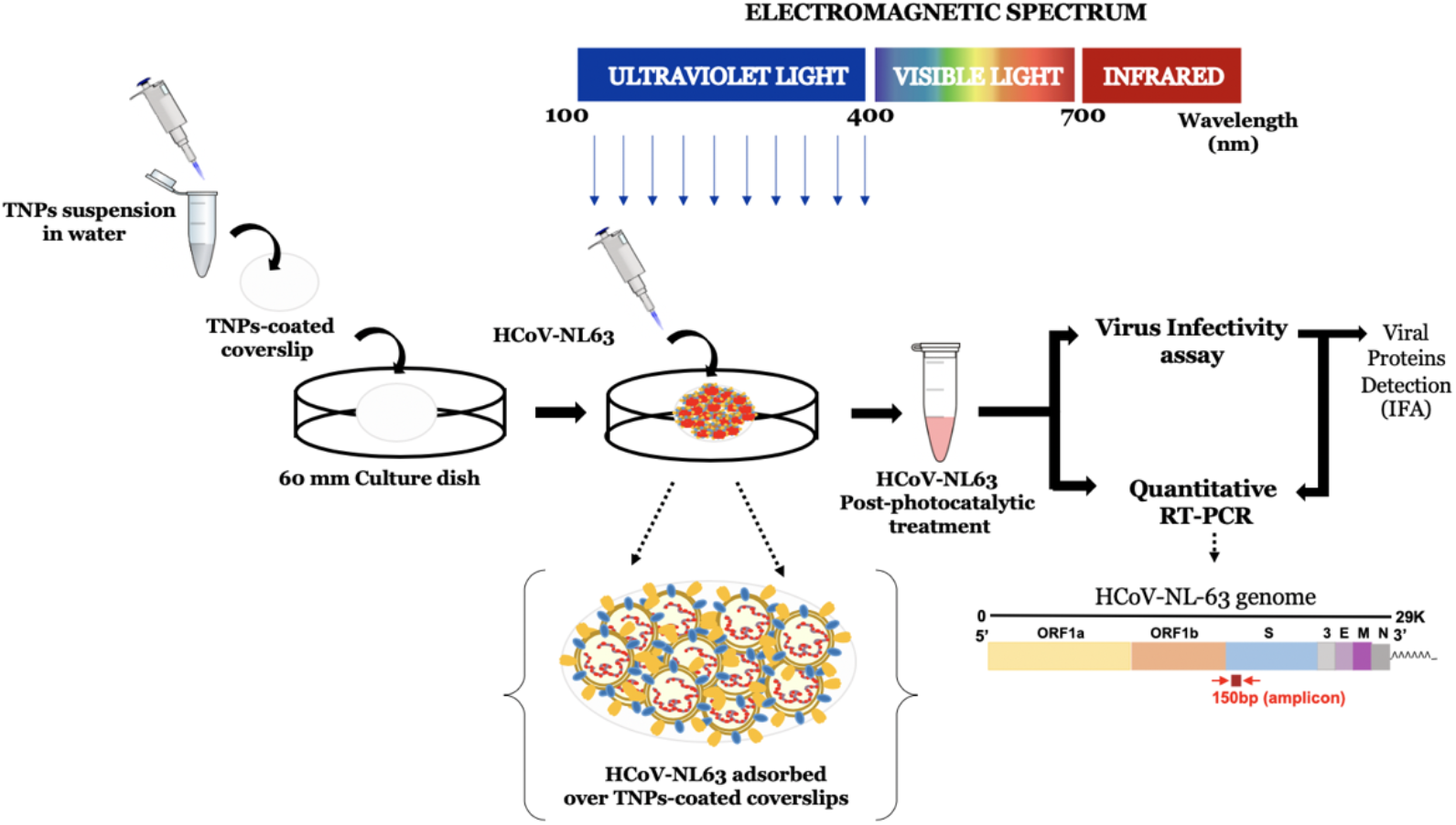
Schematic representation of the experimental set up utilized for the evaluation of photocatalytic inactivation of HCoV-NL63. TiO_2_ nanoparticles (TNPs) were prepared as a suspension in water and deposited on glass coverslips by drying at 60°C for 4-5 h. An aliquot (100 μL) with calculated virions of HCoV-NL63 was placed on the TNP-coated coverslips and exposed to UV for indicated times. The TNP/UV-treated virus was then collected to determine the viral RNA stability and infectivity via quantitative RT-PCR, infectivity and the detection of viral protein through immunofluorescence assay in HEK293L cells.

### 2.8. Immunofluorescence analysis

Cells were fixed using 3:1 methanol:acetone and stored at −80°C until used. Slides were permeabilized with 0.1% Triton X-100 for 30 min, washed (3x) and blocked (3% normal donkey serum, 0.5% BSA) for 1h at RT. Cell monolayers were washed again (3x) and incubated with rabbit anti-CoV antibody (1:200; BEI Resources, NIAID, NIH) for 1h at RT, followed by incubation with goat anti-rabbit Alexa Fluor 488 (1:5000; Molecular Probe, Carlsbad, CA), secondary antibody for 1h at RT in the dark. Finally, the nuclei were stained with TOPRO-3 (ThermoFisher, Walthman, MA). Coverslips were mounted on the glass slides using antifade and the slides were examined using Carl Zeiss LSM 780 confocal laser-scanning microscope.

### 2.9. Statistical analysis

Data presented are an average of three independent experiments and the error bars represent standard deviation. Statistical analyses were performed using Prism 8.0 software (Graphpad Inc.) and the p-values were calculated using 2-way ANOVA and the p-values are *, <0.1; and ** <0.01.

## 3. Results

### 3.1. The effect of UV light exposure on HCoV-NL63 virus stability

HCoV-NL63 virus (100 μL suspension) was applied onto the coverslip placed inside the wells of 12-well plate. The plate containing HCoV-NL63 viruses was exposed, after removing the cover, to the UV light inside a BSL-2 biological safety cabinet for 0, 1, 5, 10- and 30-min. UV-exposed virus was collected for the evaluation of virus inactivation by direct genomic RNA quantitation (qRT-PCR) and infectivity assay on HEK293L monolayer. Total RNA was extracted and quantified for genomic RNA fragmentation due to cross-linking and oxidative damage by UV exposure (Fig. 2A). HCoV-NL63 viral copies were calculated using standard curve generated on the basis of serial dilutions of known HCoV-NL63 genomic RNA. Expectedly, the calculated copies of the viral genomic RNA declined with UV-exposure (Fig. 2A). Importantly, we saw an efficient reduction in the number of intact viral genomic RNA even at 1 min exposure to the UV-light but there were still some intact copies of HCoV-NL63, at least in the region (spike protein) used for detection in our qPCR assay (Fig. 1). Increasing the exposure time to 5, 10 and 30 min, completely degraded the genomic RNA of HCoV-NL63, below the detection limit, confirming efficient inactivation of viruses with UV (Fig. 2A). Since, the qRT-PCR determines the intactness of genomic RNA, we further determined whether the UV-treatment reduced the infectivity of HCoV-NL63 virus. To this end, UV exposed HCoV-NL63 viruses were added onto HEK293L cell monolayer for 2 h to facilitate attachment and infection. Total RNA collected 48 hpi (hour post infection) was used for the detection of HCoV-NL63 genomic RNA, to measure the levels of any residual live virus in UV exposed samples (Fig. 2B). Our data showed that UV-exposure to 5-min and longer completely eliminated all infectious viruses (Fig. 2B). However, we detected some infectious viruses of HCoV-NL63, although significantly reduced, at 1 min UV exposure, suggesting 1-min exposure is insufficient in inactivating the virus completely (Fig. 2B).

**Figure 2.**
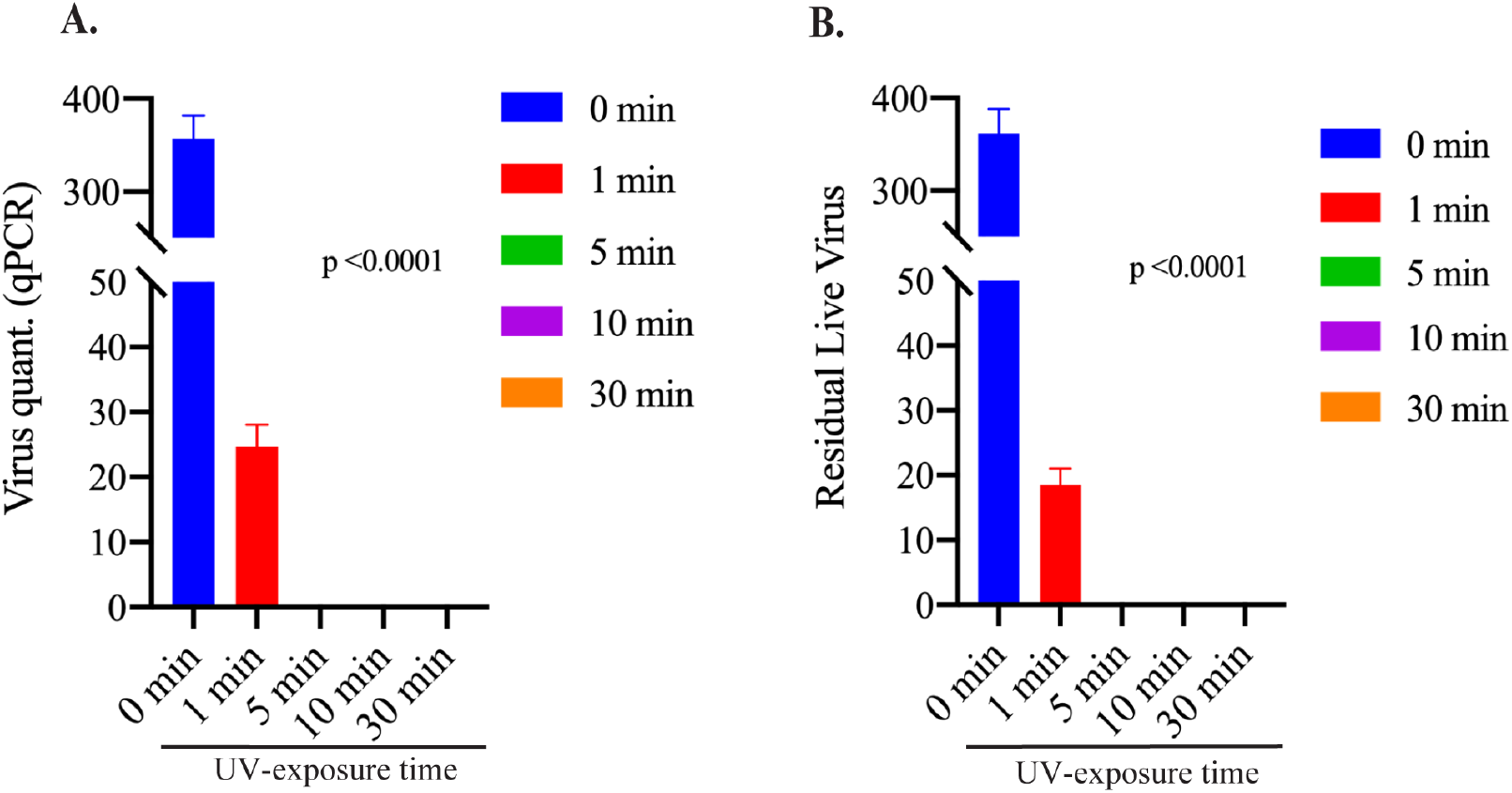
The effect of UV exposure on HCoV-NL63 RNA stability. **A.** HCoV-NL63 virus (100μL) was placed on coverslips and exposed to UV light for indicated time (0, 1, 5, 10 and 30 min). Total RNA was extracted and used for the detection viral copies by qRT-PCR. **B.** HCoV-NL63 virus (100μL) was exposed to UV light for indicated time (0, 1, 5, 10 and 30 min), collected and use for the infection of HEK293L cells. Total RNA was extracted 48h post-infection for the detection of replicated viral copies in a qRT-PCR assay. Viral copies were calculated based on a standard curve generated using the known amounts of virus obtained from BEI Resources.

### 3.2. TNP coating efficiently inactivated HCoV-NL63 even at brief exposures

Next, we sought to determine whether the photocatalytic activity of TNPs can facilitate/enhance the efficacies of virus inactivation. In order to do that, we exposed HCoV-NL63 virus with UV on the surfaces coated with TNPs, which was achieved by depositing nanosized TNPs (600 μl) onto the coverslip (2.5 cm^2^ surface area) (Fig. 1). HCoV-NL63 coronavirus (100 μL) was applied onto TNPs-coated and control coverslips (uncoated) placed inside the wells of 12-well plate and exposed to the UV light for indicated times. We quantified the number of intact HCoV-NL63 genomic RNA through RT-qPCR and infectious viral particles, as described above. Importantly, the viral copies quantified through RT-qPCR for viral genome fragmentation showed significantly reduced (almost to the background level) copies of the HCoV-NL63 from the TNP coated surface as compared to the control (uncoated surface) even at 1-min of UV-exposure (Fig. 3).

**Figure 3.**
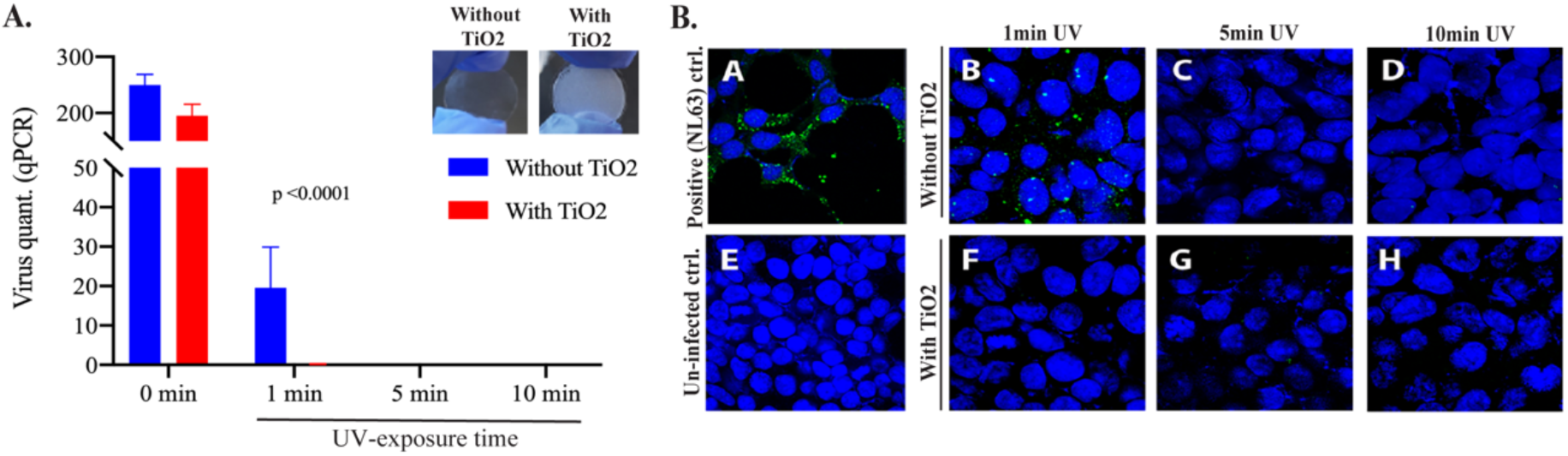
TiO_2_ nanoparticles enhanced HCoV-NL63 inactivation with no live virus in 1 min UV exposure. **(A)** HCoV-NL63 virus aliquot (100μL) was placed on glass coverslips treated with or without TiO_2_ and exposed to UV light for 1, 5- and 10min. Post-UV treatment, HCoV-NL63 was collected for viral RNA extraction. Intact viral copies were calculated based on the standard curve generated above. On top, coverslips without and with TiO2 are presented; **(B).** Post-UV treatment, HCoV-NL63 virus was collected and subjected to the infection of HEK293L monolayer. Viral protein, indicator of virus infectivity and replication, was detected by immune localization through IFA (green fluorescent dots). A) NL63 infection (positive control); B) 1 min UV light exposure without TiO_2_; C) 5 min UV light exposure without TiO_2_; D) 10 min UV light exposure without TiO_2_; E) uninfected control (negative control); F) 1 min UV light exposure with TiO_2_; G) 5 min UV light exposure with TiO_2_; H) 10 min UV light exposure with TiO_2_.

Slight variation in the number of HCoV-NL63 at 0 min UV-exposure could be because of the anti-microbial property of TNP even in the absence of UV exposure. Expectedly, the number of intact HCoV-NL63 genomic RNA was almost to the background levels at longer UV-exposure with and without TNPs (Fig. 3A). To correlate virus inactivation with infectivity, viruses recovered after UV-exposure were added onto the monolayer of HEK293L cells. The infectious viral copies were determined by localizing the protein of HCoV-NL63 through immunofluorescence assay, which is produced following replication of any residual virus. As a control of infection and replication, HCoV-NL63 was added onto HEK293L cells, and stained for viral protein to demonstrate the functionality of our assay. Detection of signals in HCoV-NL63 infected cells but not in uninfected, control confirmed the specificity of our assay (Fig. 3B, compare panel A with E). Our data showed that viruses from TNPs coated surfaces were completely inactivated even at 1 min exposure to UV-light, while surfaces without TNPs coatings had detectable levels of infectious virus, demonstrated by viral proteins (Fig. 3B, compare panels B and F). Longer exposure to UV completely inactivated the virus demonstrated by a lack of protein localization in the HEK293L infected cells (Fig. 3B, panels C, D, G, H). These data confirmed that TNPs can enhance the inactivation efficacies and brief exposure to UV-light can effectively degrade HCoV-NL63 virus rendering them non-infectious.

### 3.3. Reducing the exposure distance and relative humidity enhanced HCoV-NL63 inactivation

We further determined whether decreasing the distance of UV-source from the contaminated surface will reduce the exposure time for complete disinfection. To this end, we placed a known (calculated) amounts of HCoV-NL63 virus on a normal or TNP coated coverslips and exposed them with UV source from a distance of 10cm or 50 cm, which provided a power of 13,000μM/cm2 and 4,300μM/cm2, respectively. Expectedly, exposing the surface from a distance of 50cm (4,300μM/cm2) inactivated the virus more effectively (below the detection limit) with only limited number of detectable viral genome at 1 min exposure. Importantly, TiO2 significantly decreased the exposure time to 0.5min to achieve almost complete disinfection of the surfaces (Fig. 4A). Further decreasing the distance of exposure to 10 cm, inactivated the virus very quickly to almost background levels even in less than 0.5 min both TiO2 coated and uncoated surfaces (Fig. 4A). However, it is practically not possible and unsafe to expose surfaces with that high intensity of UV for microbial inactivation in big spaces and buildings. Therefore, we performed our assays at previously used distance (76 cm distance with 2,900μM/cm2) for our experiments determining the effects of relative humidity on HCoV-Nl63 inactivation on TNP coated surfaces. Virus placed on TNP coated and uncoated coverslips were placed under three different relative humidity and exposed to UV for indicated times. Our results showed that increase in the relative humidity enhances HCoV-NL63 inactivation even on normal surfaces (Fig. 4B). Importantly, TNP coating enhanced the inactivation of HCoV-NL63 virus at all three tested relative humidity (Fig. 4B). This confirmed that surface inactivation of viruses can be achieved with shorter UV exposure on TNP coated surfaces and the humidity enhances the photocatalytic activity of TiO2.

**Figure 4.**
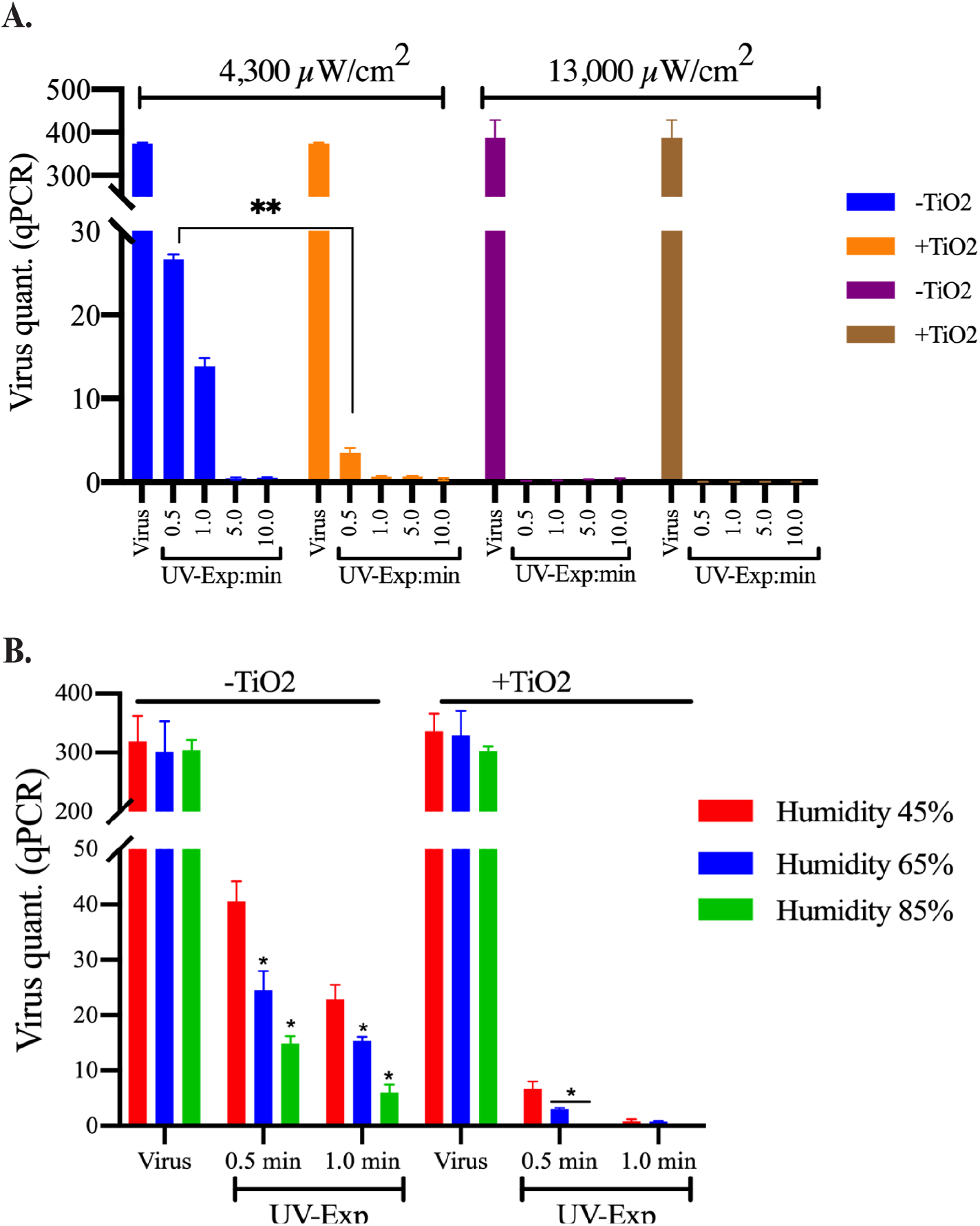
UV energy and humidity affected TNP mediated virus inactivation. **(A)** HCoV-NL63 virus aliquot (100 μL) was placed on glass coverslips treated with or without TiO_2_ and exposed to UV light for 0.5, 1, 5, and 10-min from a distance of 50cm or 10cm to achieve 4,300μM/cm2 and 13,000μM/cm2 energy levels. Post-UV treatment, HCoV-NL63 was collected for viral RNA extraction. Intact viral copies were calculated based on the standard curve generated above. **(B).** HCoV-NL63 virus aliquot (100μL) placed on TNP coated or control (uncoated) coverslips were exposed to UV-lights for 0.5 and 1.0 min under 45%, 65% and 85% relative humidity. Virus from UV-untreated surfaces were used as controls. Post-UV treatment, viral genomic RNA was extracted and subjected for quantitation of genomic RNA in qPCR.

## 4. Discussion

Lack of anti-COVID-19 vaccine and highly contagious nature of SARS-CoV-2 highlights the need for environmental control of viral spread. Stability of virus in different environmental conditions [22] and ability to spread far distances made this virus exceptionally successful in dissemination within the human population [23,24]. SARS-CoV-2 appears to be highly stable on smooth surfaces for up to 72h and in a wide range of pH conditions (pH 3-10) at room temperature (22°C) [24]. Virus can spread on surfaces by direct contact or aerosol droplets. The small size droplets (10μm) can become airborne and circulate in the air for an extended period of time carrying the infectious virus to far distances [25]. They may be dispersed widely by air contaminating the environment far distant from the patient. Therefore, decontaminating surfaces is essential to combat COVID-19 spread.

Nanosized TiO_2_/TNPs have photocatalytic properties, which are shown to control microbial growth [15,26]. The mechanism of antimicrobial activity is based on the excitation of the electron/e^-^ from the valence band to the conduction band and the generation of the “electron-hole (e^-^h+)” [27]. The free electron can contribute to production of the reactive oxygen species (ROS) including O_2_^-^ and OH^-^ radicals [28]. These ROS can react in the solution producing highly potent anti-microbial H_2_O_2_. The TNPs can be used in suspension, liquids or immobilized on surfaces, also referred as a “self-cleaning” surfaces [29]. This feature of TNPs could provide a microbial control in addition to conventional disinfecting products. TNP’s anti-viral effect has been shown against influenza virus, Newcastle virus, hepatitis B virus and herpesviruses [30–33]. TiO_2_ was proposed to have antiviral effect, targeting both DNA and RNA viruses, airborne and blood borne pathogens. The photocatalytic titanium apatite filter was earlier shown to inactivate SARS coronavirus up to 99.99% after 6 h exposure [34]. Our assay to use TiO_2_ coating to efficiently inactivate human coronavirus with UV light through very brief exposure highlights the importance of TNP coatings on publicly used surfaces. Although, UV by itself are used for viral inactivation including the trains of public transport systems but the efficacies are not well established. Since our assays showed augmentation of virus inactivation, HCoV-NL63 by the TNPs coatings under UV exposure, we propose that TNP coatings can enhance virus inactivation including SARS-CoV-2 on surfaces most frequently exposed with COVID-19 patients. HCoV-NL63 and SARS-CoV-2 both cause acute respiratory diseases and have similar virion structure (HCoV) [18,35], we speculate that these inactivation parameters will be same for the both viruses.

Various factors are shown to affect virus resistance to UV light exposure, including UV dose, humidity and the surface material [36,37]. It was reported that the virus survival decreases with an increase in the UV energy [38]. Recently, Buonnano et al have demonstrated that alpha and beta coronaviruses are susceptible to the low dose of the UV light exposure, where 90% of virus inactivation was achieved in 8 minutes [39]. The UV light susceptibility of SARS-CoV-2 was also demonstrated, confirming the virucidal effect of the UV light [40]. In both studies, the effect of the higher UV light intensity was shown to have more virucidal effect, as expected [39,40]. In line with these data, we have shown the beta coronavirus is susceptible to the UV radiation and that this effect is directly proportional to the UV-light intensity. The most interesting observation is that TiO_2_ can facilitate virucidal effect of the low intensity UV light, which could have important practical application. As TiO2 can maintain antiviral efficacy of the UV light at the lower intensity, it will help in reducing the potential harmful effect of radiation exposure [41].

Interestingly, TiO_2_ nanoparticles effectively inactivated HCoV-NL63 under all three tested relative humidity conditions, which are within the standard for the indoor conditions (30-60%) [42]. We have shown that TiO_2_ nanoparticles retain virucidal efficacy even at very high humid condition (85% relative humidity), confirming a broader use of TNPs coatings on outdoor surfaces. The environmental humidity was suggested to play a role in the airborne virus spread [43–45]. It is postulated that humid environment contributes to the size of droplets and the stability of the virions [25,46]. A prolonged survival of the airborne viruses was demonstrated in the lower humid conditions [44,47]. Study by Wu et al have demonstrated a negative correlation between COVID-19 cases and humidity levels [48]. Study published by Matson et al [49], showed that lower humidity combined with lower temperature prolonged the half-life of virus in nasal mucosa. Matson et al also stated that virus is maintained in the climate controlled environment with relative humidity 40%, for 24 hours [49].

A presumed model to understand the underlying mechanism involved in the photocatalytic inactivation of HCoV-NL63 has been described in Fig. 5. In the presence of UV radiation, TiO_2_ nanoparticles act as photocatalyst due to their specific energy structure. UV-irradiation of nanosized TiO_2_ results in the excitation of electron leading to the formation of a specific “electron-hole (e^-^-h^+^)” pair and generation of reactive oxygen species (ROS). The electron holes (h^+^) catalyze oxidation processes and convert water/hydroxide molecules to peroxide/hydroxyl radicals, whereas electrons (e^-^) induce reduction reactions and react with molecular oxygen to generate superoxide radicals. These resulting ROS have the ability to inactivate the virus through oxidative damage.

**Figure 5.**
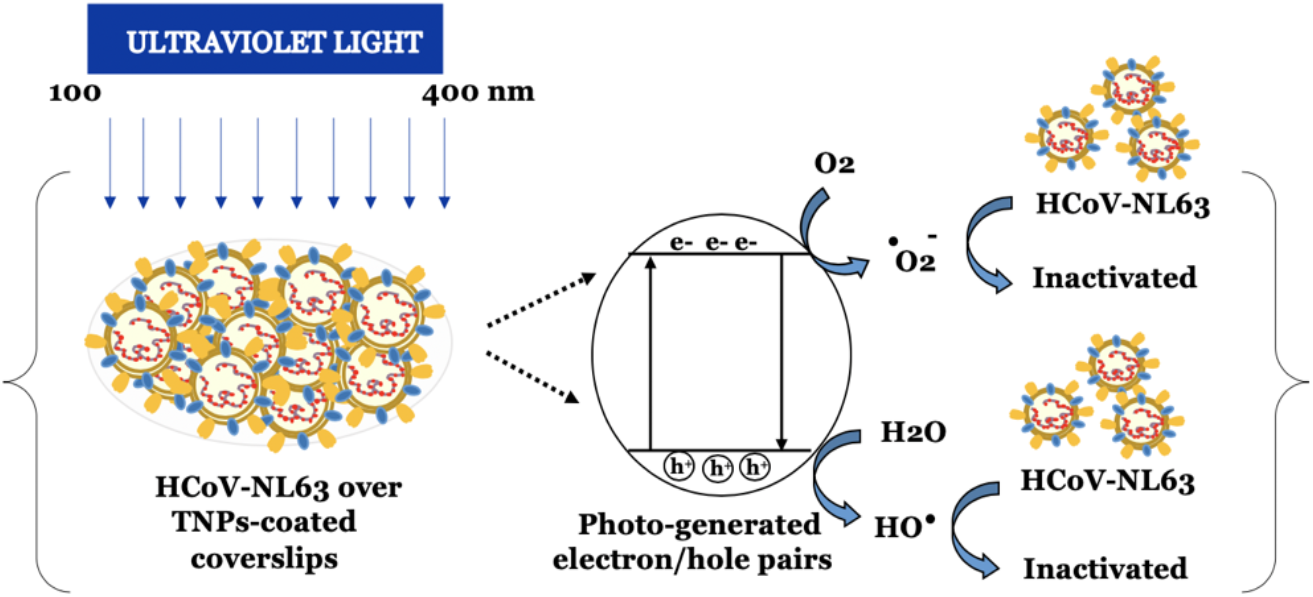
Schematic illustration of photocatalytic inactivation of HCoV-NL63 principle. Irradiation with UV light generates specific “electron-hole (e^-^-h^+^)” pair, and reactive oxygen species (ROS), including hydroxyl and superoxide radicals on the surface of TNPs. These ROS have strong oxidative ability, which can inactivate the virus due to oxidative damage.

Since, TiO_2_ is non-toxic and used as a food additives [50], although at low quantities, TNPs long-term effects on human health has not been evaluated. Therefore, a well-controlled toxicology studies along with approvals from the environmental protection agency and associated regulatory bodies may be needed, before using the TNPs on commonly used surfaces. TNPs have many advantages as compared to conventional germicidal products including its low cost and chemical stability [51]. Additionally, they do not require multiple applications as they could be coated single time on various surfaces [52,53]. In fairness, TNPs can be applied as a paint on the surfaces in the form of immobilized thin coatings on high risk facilities including train stations and other public areas to reduce the surface contamination and any possible exposures. It is obvious advantage of TNPs that they have a broad range of anti-microbial efficacy including virucidal, bactericidal and fungicidal [15]. This is especially beneficial in the areas with high risk of exposure to biohazardous material, such as hospitals. Additionally, this may be useful in hospital settings including the ICUs handling COVID-19 patients, where TNP coatings of the hospital floors can provide an effective way of inactivating viruses and surface contamination of SARS-CoV-2. In conclusion, we highlight the effectiveness of UV light-induced virucidal activity of TNPs in controlling coronavirus spread, which can help create a sanitary environment.

## Author Contributions

“Conceptualization, R.S and S.C.V.; methodology, S.K., T.U., N.D., R.S. and S.C.V.; formal analysis, S.K., T.U., and S.C.V.; investigation, S.K., T.U., and S.C.V.; resources, R.S and S.C.V.; data curation, S.K.; writing—original draft preparation, S.K.; writing— review and editing, S.K., T.U., and S.C.V.; supervision, R.S and S.C.V.; project administration, S.C.V and R.S. All authors have read and agreed to the published version of the manuscript.

## Acknowledgments

Human Coronavirus, NL63 (HCoV-NL63) was obtained through BEI Resources, NIAID, NIH: Human Coronavirus, NL63, NR-470. Serum used in the assay was obtained through BEI Resources, NIAID, NIH: Rabbit Sera Control Panels, Polyclonal Anti-SARS-CoV Spike Protein, NR-4569.

## References

1. CENTER, C.R. COVID-19 Dashboard by the Center for Systems Science and Engineering (CSSE) at Johns Hopkins University (JHU). Availabe online: https://coronavirus.jhu.edu/map.html (accessed on

2. Organization, W.H. Coronavirus disease 2019 (COVID-19): situation report, 30. 2020.

3. Control, C.f.D.; Prevention. Provisional death counts for coronavirus disease (covid-19). 2020.

4. Shao, L.; Li, X.; Zhou, Y.; Yu, Y.; Liu, Y.; Liu, M.; Zhang, R.; Zhang, H.; Wang, X.; Zhou, F. Novel Insights Into Illness Progression and Risk Profiles for Mortality in Non-survivors of COVID-19. Front Med (Lausanne) 2020, 7, 246, doi:10.3389/fmed.2020.00246.

5. Burns, E.; Kakara, R. Deaths from Falls Among Persons Aged >/=65 Years - United States, 2007-2016. MMWR Morb Mortal Wkly Rep 2018, 67, 509–514, doi:10.15585/mmwr.mm6718a1.

6. Guan, W.-j.; Ni, Z.-y.; Hu, Y.; Liang, W.-h.; Ou, C.-q.; He, J.-x.; Liu, L.; Shan, H.; Lei, C.-l.; Hui, D.S. Clinical characteristics of coronavirus disease 2019 in China. New England journal of medicine 2020, 382, 1708–1720.

7. Van Doremalen, N.; Bushmaker, T.; Morris, D.; Holbrook, M.; Gamble, A.; Williamson, B. & Lloyd-Smith, JO (2020). Aerosol and surface stability of SARS-CoV-2 as compared with SARS-CoV-1. New England journal of medicine.

8. Huang, Y.; Chen, S.; Yang, Z.; Guan, W.; Liu, D.; Lin, Z.; Zhang, Y.; Xu, Z.; Liu, X.; Li, Y. SARS-CoV-2 viral load in clinical samples of critically ill patients. American journal of respiratory and critical care medicine 2020.

9. Morawska, L. Droplet fate in indoor environments, or can we prevent the spread of infection? In Proceedings of Proceedings of Indoor Air 2005: the 10th International Conference on Indoor Air Quality and Climate; pp. 9–23.

10. Wang, J.; Du, G. COVID-19 may transmit through aerosol. Irish Journal of Medical Science (1971-) 2020, 1–2.

11. Organization, W.H. Coronavirus disease (COVID-19) advice for the public. 2020.

12. Ong, S.W.X.; Tan, Y.K.; Chia, P.Y.; Lee, T.H.; Ng, O.T.; Wong, M.S.Y.; Marimuthu, K. Air, surface environmental, and personal protective equipment contamination by severe acute respiratory syndrome coronavirus 2 (SARS-CoV-2) from a symptomatic patient. Jama 2020.

13. Liu, Y.; Ning, Z.; Chen, Y.; Guo, M.; Liu, Y.; Gali, N.K.; Sun, L.; Duan, Y.; Cai, J.; Westerdahl, D., et al. Aerodynamic analysis of SARS-CoV-2 in two Wuhan hospitals. Nature 2020, 582, 557-560, doi:10.1038/s41586-020-2271-3.

14. Cai, J.; Sun, W.; Huang, J.; Gamber, M.; Wu, J.; He, G. Indirect Virus Transmission in Cluster of COVID-19 Cases, Wenzhou, China, 2020. Emerging Infectious Diseases 2020, 26.

15. Bogdan, J.; Zarzyńska, J.; Pławińska-Czarnak, J. Comparison of infectious agents susceptibility to photocatalytic effects of nanosized titanium and zinc oxides: a practical approach. Nanoscale research letters 2015, 10, 1–15.

16. Nakano, R.; Ishiguro, H.; Yao, Y.; Kajioka, J.; Fujishima, A.; Sunada, K.; Minoshima, M.; Hashimoto, K.; Kubota, Y. Photocatalytic inactivation of influenza virus by titanium dioxide thin film. Photochemical & Photobiological Sciences 2012, 11, 1293–1298.

17. Mazurkova, N.; Spitsyna, Y.E.; Shikina, N.; Ismagilov, Z.; Zagrebel’nyi, S.; Ryabchikova, E. Interaction of titanium dioxide nanoparticles with influenza virus. Nanotechnologies in Russia 2010, 5, 417–420.

18. Abdul-Rasool, S.; Fielding, B.C. Understanding human coronavirus HCoV-NL63. The open virology journal 2010, 4, 76.

19. ATCC. Human coronavirus 229E (ATCC^®^ VR-740™) Availabe online: https://www.lgcstandards-atcc.org/products/all/VR-740.aspx?geo_country=ru (accessed on

20. Control, C.f.D.; Prevention. INDOOR ENVIRONMENTAL QUALITY. Availabe online: (accessed on

21. Current, R. Annual Average Relative Humidity by US State. Availabe online: https://www.currentresults.com/Weather/US/annual-average-humidity-by-state.php (accessed on

22. Desai, A.N.; Patel, P. Stopping the Spread of COVID-19. Jama 2020, 323, 1516–1516.

23. Casanova, L.M.; Jeon, S.; Rutala, W.A.; Weber, D.J.; Sobsey, M.D. Effects of air temperature and relative humidity on coronavirus survival on surfaces. Appl. Environ. Microbiol. 2010, 76, 2712–2717.

24. Chin, A.; Chu, J.; Perera, M. Correspondence. Stability of SARS-CoV-2 in different environmental conditions. Lancet Microbe 2020, 1, 10, doi:https://doi.org/10.1016/S2666-5247(20)30003-3.

25. Gralton, J.; Tovey, E.; McLaws, M.-L.; Rawlinson, W.D. The role of particle size in aerosolised pathogen transmission: a review. Journal of Infection 2011, 62, 1–13.

26. Tatlidil, Ì.; Sökmen, M.; Breen, C.; Clegg, F.; Buruk, C.K.; Bacaksız, E. Degradation of Candida albicans on TiO 2 and Ag-TiO 2 thin films prepared by sol–gel and nanosuspensions. Journal of sol-gel science and technology 2011, 60, 23.

27. Panayotov, D.A.; Burrows, S.P.; Morris, J.R. Photooxidation mechanism of methanol on rutile TiO2 nanoparticles. The Journal of Physical Chemistry C 2012, 116, 6623–6635.

28. Vatansever, F.; de Melo, W.C.; Avci, P.; Vecchio, D.; Sadasivam, M.; Gupta, A.; Chandran, R.; Karimi, M.; Parizotto, N.A.; Yin, R. Antimicrobial strategies centered around reactive oxygen species–bactericidal antibiotics, photodynamic therapy, and beyond. FEMS microbiology reviews 2013, 37, 955–989.

29. Yates, H.; Brook, L.; Ditta, I.; Evans, P.; Foster, H.; Sheel, D.; Steele, A. Photo-induced self-cleaning and biocidal behaviour of titania and copper oxide multilayers. Journal of photochemistry and photobiology A: Chemistry 2008, 197, 197–205.

30. Zan, L.; Fa, W.; Peng, T.; Gong, Z.-k. Photocatalysis effect of nanometer TiO2 and TiO2-coated ceramic plate on Hepatitis B virus. Journal of Photochemistry and Photobiology B: Biology 2007, 86, 165–169.

31. van der Molen, R.G.; Garssen, J.; de Klerk, A.; Claas, F.H.; Norval, M.; van Loveren, H.; Koerten, H.K.; Mommaas, A.M. Application of a systemic herpes simplex virus type 1 infection in the rat as a tool for sunscreen photoimmunoprotection studies. Photochemical & Photobiological Sciences 2002, 1, 592–596.

32. Monmaturapoj, N.; Sri-on, A.; Klinsukhon, W.; Boonnak, K.; Prahsarn, C. Antiviral activity of multifunctional composite based on TiO2-modified hydroxyapatite. Materials Science and Engineering: C 2018, 92, 96–102.

33. Akhtar, S.; Shahzad, K.; Mushtaq, S.; Ali, I.; Rafe, M.H.; Fazal-ul-Karim, S.M. Antibacterial and antiviral potential of colloidal Titanium dioxide (TiO2) nanoparticles suitable for biological applications. Materials Research Express 2019, 6, 105409.

34. Han, W.; Zhang, B.; Cao, W.; Yang, D.; Taira, I.; Okamoto, Y.; Arai, J.; Yan, X. The inactivation effect of photocatalytic titanium apatite filter on SARS virus. Sheng wu hua xue yu sheng wu wu li jin zhan 2004, 31, 982–985.

35. Xia, S.; Liu, M.; Wang, C.; Xu, W.; Lan, Q.; Feng, S.; Qi, F.; Bao, L.; Du, L.; Liu, S. Inhibition of SARS-CoV-2 (previously 2019-nCoV) infection by a highly potent pan-coronavirus fusion inhibitor targeting its spike protein that harbors a high capacity to mediate membrane fusion. Cell research 2020, 30, 343–355.

36. McDevitt, J.J.; Rudnick, S.N.; Radonovich, L.J. Aerosol susceptibility of influenza virus to UV-C light. Applied and environmental microbiology 2012, 78, 1666–1669.

37. Van Doremalen, N.; Bushmaker, T.; Morris, D.H.; Holbrook, M.G.; Gamble, A.; Williamson, B.N.; Tamin, A.; Harcourt, J.L.; Thornburg, N.J.; Gerber, S.I. Aerosol and surface stability of SARS-CoV-2 as compared with SARS-CoV-1. New England journal of medicine 2020, 382, 1564–1567.

38. Tseng, C.-C.; Li, C.-S. Inactivation of virus-containing aerosols by ultraviolet germicidal irradiation. Aerosol Science and Technology 2005, 39, 1136–1142.

39. Buonanno, M.; Welch, D.; Shuryak, I.; Brenner, D.J. Far-UVC light (222 nm) efficiently and safely inactivates airborne human coronaviruses. Scientific Reports 2020, 10, 1–8.

40. Bianco, A.; Biasin, M.; Pareschi, G.; Cavalleri, A.; Cavatorta, C.; Fenizia, F.; Galli, P.; Lessio, L.; Lualdi, M.; Redaelli, E. UV-C irradiation is highly effective in inactivating and inhibiting SARS-CoV-2 replication. Inactivating and Inhibiting SARS-CoV-2 Replication (June 5, 2020) 2020.

41. Wilson, B.D.; Moon, S.; Armstrong, F. Comprehensive review of ultraviolet radiation and the current status on sunscreens. The Journal of clinical and aesthetic dermatology 2012, 5, 18.

42. Arena, L.B.; Karagiozis, A.; Mantha, P. Monitoring of Internal Moisture Loads in Residential Buildings—Research Findings in Three Different Climate Zones. In Proceedings of Buildings XI: Thermal Performance of Exterior Envelopes of Whole Buildings International Conference, Clearwater Beach, FL: ASHRAE. Retrieved from http://web.ornl.gov/sci/buildings/2012/2010%20B11%20papers/75_Arena.pdf.

43. Yang, W.; Marr, L.C. Dynamics of airborne influenza A viruses indoors and dependence on humidity. PloS one 2011, 6, e21481.

44. Chan, K.-H.; Peiris, J.M.; Lam, S.; Poon, L.; Yuen, K.; Seto, W.H. The effects of temperature and relative humidity on the viability of the SARS coronavirus. Advances in virology 2011, 2011.

45. Berry, A.; Fillaux, J.; Martin-Blondel, G.; Boissier, J.; Iriart, X.; Marchou, B.; Magnaval, J.F.; Delobel, P. Evidence for a permanent presence of schistosomiasis in Corsica, France, 2015. Eurosurveillance 2016, 21, 30100.

46. Sooryanarain, H.; Elankumaran, S. Environmental role in influenza virus outbreaks. Annu. Rev. Anim. Biosci. 2015, 3, 347–373.

47. Weber, T.P.; Stilianakis, N.I. Inactivation of influenza A viruses in the environment and modes of transmission: a critical review. Journal of Infection 2008, 57, 361–373.

48. Wu, Y.; Jing, W.; Liu, J.; Ma, Q.; Yuan, J.; Wang, Y.; Du, M.; Liu, M. Effects of temperature and humidity on the daily new cases and new deaths of COVID-19 in 166 countries. Science of the Total Environment 2020, 139051.

49. Matson, M.J.; Yinda, C.K.; Seifert, S.N.; Bushmaker, T.; Fischer, R.J.; van Doremalen, N.; Lloyd-Smith, J.O.; Munster, V.J. Early Release-Effect of Environmental Conditions on SARS-CoV-2 Stability in Human Nasal Mucus and Sputum.

50. Weir, A.; Westerhoff, P.; Fabricius, L.; Hristovski, K.; Von Goetz, N. Titanium dioxide nanoparticles in food and personal care products. Environmental science & technology 2012, 46, 2242–2250.

51. Bosc, F.; Lacroix-Desmazes, P.; Ayral, A. TiO2 anatase-based membranes with hierarchical porosity and photocatalytic properties. J Colloid Interface Sci 2006, 304, 545–548, doi:10.1016/j.jcis.2006.09.064.

52. Visai, L.; De Nardo, L.; Punta, C.; Melone, L.; Cigada, A.; Imbriani, M.; Arciola, C.R. Titanium oxide antibacterial surfaces in biomedical devices. The International journal of artificial organs 2011, 34, 929–946.

53. De Jong, B.; Meeder, A.; Koekkoek, K.; Schouten, M.; Westers, P.; Van Zanten, A. Pre–post evaluation of effects of a titanium dioxide coating on environmental contamination of an intensive care unit: the TITANIC study. Journal of Hospital Infection 2018, 99, 256–262.

